# Coinfinder: Detecting Significant Associations and Dissociations in Pangenomes

**DOI:** 10.1101/859371

**Authors:** Fiona Jane Whelan, Martin Rusilowicz, James Oscar McInerney

## Abstract

2.

The accessory genes of prokaryote and eukaryote pangenomes accumulate by horizontal gene transfer, differential gene loss, and the effects of selection and drift. We have developed Coinfinder, a software program that assesses whether sets of homologous genes (gene families) in pangenomes associate or dissociate with each other (i.e. are “coincident”) more often than would be expected by chance. Coinfinder employs a user-supplied phylogenetic tree in order to assess the lineage-dependence (i.e. the phylogenetic distribution) of each accessory gene, allowing Coinfinder to focus on coincident gene pairs whose joint presence is not simply because they happened to appear in the same clade, but rather that they tend to appear together more often than expected across the phylogeny. Coinfinder is implemented in C++, Python3, and R and is freely available under the GPU license from https://github.com/fwhelan/coinfinder.

**Impact statement:** Coinfinder identifies genes that co-occur (associate) or avoid (dissociate) with each other across the accessory genomes of a pangenome of interest. Genes that associate or dissociate more often than expected by chance, suggests that those genes have a connection (attraction or repulsion) that is interesting to explore. Identification of these groups of genes will further the field’s understanding of the importance of accessory genes. Coinfinder is a freely available, open-source software which can identify gene patterns locally on a personal computer in a matter of hours.

**Data summary:**

1. Coinfinder is freely available at https://github.com/fwhelan/coinfinder.
2. A list of the Identifiers of the genomes used within as well as all input/output files are available at https://github.com/fwhelan/coinfinder-manuscript.

**The authors confirm all supporting data, code and protocols have been provided within the article or through supplementary data files.**

## 5. Introduction

Pangenomes consist of core genes, common across all strains of a species, and accessory genes that are present in some but not all strains (1). Accessory genes by definition are not essential to the existence of a species, therefore it remains somewhat unclear why accessory genes exist, and what influences the content of these accessory genomes. It is likely that some genes co-occur, or associate, because they positively influence each other’s fitness in a particular, or set of, host genomes. Similarly, we expect some genes to avoid, or dissociate with one another because their co-occurrence produces a negative fitness effect. We expect that genes whose products function together in a biochemical pathway, or that can combine to form a useful heteromeric protein complex, will appear together in the same genome more often than their observed frequency in the dataset would predict. For example, MYD88 consistently co-occurs with the genetic components of the MYD88-dependent TLR-signalling pathway in vertebrate species (2). In contrast, genes that produce a toxic by-product when they are expressed in the same cell, or that perform the same function and therefore induce functional redundancy, are expected to appear together less often than their observed frequency in the dataset would predict. This is seen, for example, with siderophore biosynthetic gene clusters in *Salinispora spp.* where an isolate either has one iron-chelating siderophore or a different non-homologous system, but never both (3). As a first step towards understanding these kinds of gene-to-gene interactions in the accessory pangenome, it is useful to identify genes that appear together or that avoid one another significantly more often than would be expected by chance.

Previously established methodology can identify various forms of co-occurrence patterns in prokaryotes. For example, many tools (e.g. (4)(5)(6)) and tool comparisons (7) are available for the identification of species-species co-occurrence patterns in microbial communities. For example, the program SparCC identifies correlations in compositional data, including species presence-absence patterns within microbial communities (8). Other tools, such as NetShift (9), find differences in species association networks of microbial communities across datasets (e.g. healthy versus diseased states). Similarly, methods have been established to identify associations between genotypic and phenotypic traits in pangenomes (i.e. gene-trait co-occurrence). Usually called pangenome genome-wide association studies (pan-GWAS), tools such as bugwas (10) and Scoary (11) compare components of the pangenome to a user-provided list of phenotypic traits. New methods such as SpydrPick (12) identify Single Nucleotide Polymorphisms (SNP)-SNP co-occurrence patterns by comparing SNPs in multiple sequence alignments of proteins in microbial population genomic datasets.

A few approaches have focussed on gene-gene co-occurrence. Pantagruel (13) uses gene- and species-trees to identify genes which have similar patterns of gain and loss in a pangenome to define co-evolved gene modules. Similarly, CoPAP (14) searches for correlated patterns of gene gain and loss across a species tree to find co-evolutionary interactions of Clustered Orthologous Groups (COGs). While conceptually similar to Coinfinder, these methodologies are based on phyletic patterns; further, the dissociation of genes isn’t considered by either method. The most similar method to Coinfinder in concept is the identification of correlogs and anti-correlogs, genes which favour or dis-favour co-occurrence within a genome, by Kim and Price (15). However, this method was not packaged into publicly available software and was not coupled with the pangenome concept, instead focusing on global patterns of gene associations across the bacterial Domain.

Here, we present Coinfinder, a command line software program that identifies coincident (associating or dissociating) genes across a set of input genomes. Coinfinder can run in any Unix environment using a user-specified number of processing cores. Coinfinder can be used to investigate the structure of strain- or species-pangenomes and is not restricted to prokaryote or eukaryote genomic input.

## 6. Theory and Implementation

### 6.1 Input

Coinfinder accepts genome content data in one of two formats: (a) the gene_presence_absence.csv output from Roary (16); or (b) as a tab-delimited list of the genes present in each strain. If option (b) is used, genes should be clustered into orthologous groups/gene clusters prior to using Coinfinder (for example, using BLAST (17) and a clustering algorithm, such as MCL (18)(19). Additionally, Coinfinder requires a Newick-formatted phylogeny of the genomes in the dataset. We suggest that this phylogeny can be constructed using the core genes from the input genomes as produced using programs such as Roary, or using ribosomal RNA genes, or a similar approach (20).

### 6.2 Identifying coincident genes

For each set of genes in the input genomes, Coinfinder examines the presence/absence pattern of the gene pair to determine if they represent a coincident relationship; i.e. if *gene i* and *gene j* are observed together or apart in the input genomes more often than would be expected by chance.

As a pre-processing step, the input gene set is culled for high- and low-abundance genes. Genes present in every genome (i.e. core genes) are removed as they cannot statistically associate or dissociate (i.e. be coincident with) another gene more or less often than expected. Similarly, genes whose presence is constrained to a small number of genomes will not produce significant associations, therefore low-abundance genes can be removed from the input at a user-determined cutoff. Coinfinder’s default is to remove any gene present in less than 5% of the input genomes.

Coinfinder has two modes for identifying coincident relationships: association and dissociation. When testing for gene associations, Coinfinder evaluates whether *gene i* and *gene j* of a given gene pair are observed together in the input genomes more often than would be expected by chance. More formally, for a set of genomes *N*, we define the probability of observing *gene i* as:

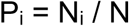

where N_i_ is the number of occurrences of *gene i* in the dataset. The expected rate of association, E_A_, of *gene i* with *gene j*, is then defined as:

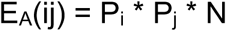

and the observed rate of association, O_A_, as:

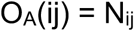

where N_ij_ is the number of times *gene i* and *gene j* are present within the same genome.

When testing gene dissociation, Coinfinder evaluates whether *gene i* and *gene j* of a given gene pair are observed separately in the input genomes more often than would be expected by chance. Formally, the expected rate of dissociation, E_D_, is defined as:

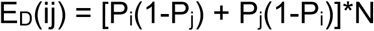

and the observed rate of dissociation, O_D_, as:

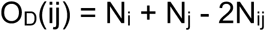

In each mode, Coinfinder’s default behaviour is to use a Bonferroni-corrected binomial exact test statistic (adapted from https://github.com/chrchang/stats) of the expected and observed rates to evaluate whether each gene pair are significantly coincident with each other.

Coincident genes that share an evolutionary history are more likely to have indirect correlations with each other. For example, if two genes are found to associate and each is observed only within a particular clade, the most parsimonious explanation for the observation is that the last common ancestor of the clade obtained both genes at the same evolutionary step. These two genes may, or may not, have a functional relationship with one another, and are of potential interest. However, non-monophyletic – or lineage-independent – genes that are dispersed throughout a phylogeny and are found to be significantly coincident are more likely to have a direct relationship with each other – their patchy phylogenetic distribution, combined with their statistically significant rate of association is *prima facie* evidence that they interact in some way. Thus, Coinfinder focuses on identifying coincident relationships between lineage-independent accessory genes. To do this, Coinfinder uses a previously established phylogenetic measure of binary traits (D, as coded into the R function phylo.d; (21)) to determine the lineage-dependence of each coincident gene. D is a measure of phylogenetic signal strength of a binary trait, which quantifies the amount of dispersion of the trait – here, the presence of a gene – over a phylogenetic tree (21).

### 6.3 Output

Coinfinder visualizes the results of its analysis in two ways. First, Coinfinder produces a network in which each node is a gene family and each edge is a statement of significant gene association (corrected for lineage effects) or significant gene dissociation. The size of a node is proportional to the gene’s D value. Second, Coinfinder generates a presence-absence heatmap, indicating the presence of coincident genes in the context of the input phylogeny. The genes in the heatmap are ordered by D value (from most lineage-independent to least) and are coloured according to coincident patterns.

Coinfinder produces a number of output files, with the default prefix of *coincident_*, as described in Table 1. Examples of the network and heatmap outputs of Coinfinder are shown in Figure 1.

**Table 1:** Description of Coinfinder output files.

| **Suffix** | **File description** |
| --- | --- |
| _pairs.tsv | Tab-delimited list of significant coincident gene pairs |
| _nodes.tsv | Node list of all unique coincident genes and their D value |
| _edges.tsv | Edge list of significant gene-gene pairs and the associated p-value |
| _network.gexf | GEXF (Graph Exchange XML Format) v1.2 formatted network file. Nodes are coloured by connected component (i.e. coincident gene set) and sized by D value; edge thickness is proportional to the p-value of the coincident relationship between any two connected genes |
| _components.tsv | Tab-delimited list of all connected components within the gene-gene coincident network |
| _heatmap[0-X].pdf | Heatmap images (R, ggplot2 (30), ggtree (31)) of the presence-absence patterns of coincident components across input genomes. The heatmap is split across multiple files when needed for ease of visibility |

**Figure 1:**
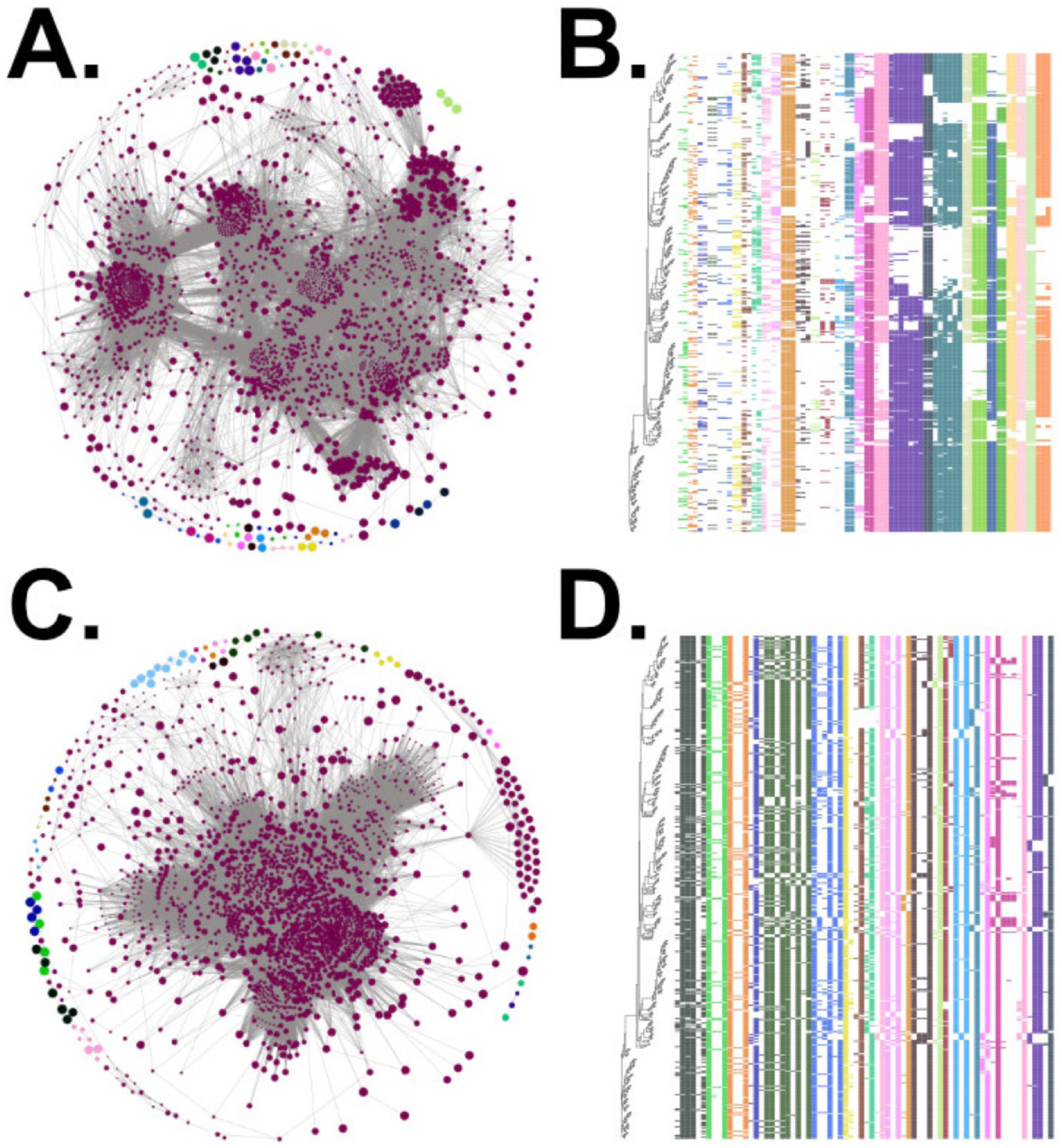
Example of Coinfinder output. The network (**A,C**) and heatmap (**B,D**) outputs from Coinfinder executed on 534 *Streptococcus pneumoniae* genomes. **A,C**. The resultant gene association (**A**) and dissociation (**C**) networks. Each gene (node) is connected to (edge) another gene if they statistically associate/dissociate with each other in the pangenome. Nodes are coloured by connected component (i.e. coincident gene sets) and the colours correspond to those used in the heatmap outputs. The network file Coinfinder generates includes all node and edge colouring; Gephi (29) was used to apply the Fruchterman Reingold layout. **B,D.** A portion of the heatmaps of the presence/absence patterns of the associating (**B**) and dissociating (**D**) gene sets. Similar to the network, each set of coincident genes are co-coloured. Genes are displayed in relation to the input core gene phylogeny. Here the phylogeny tip and gene cluster labels have been removed from the output for clarity. Additionally, the largest connected component in the network (wine colour) has been omitted from the heatmap for ease of display.

## 7. Results

As an example, Coinfinder was executed using 534 *Streptococcus pneumoniae* genomes as input, a subset of the Global Pneumococcal Sequencing Project (GPS; https://www.pneumogen.net/gps/) whose open reading frames (ORFs) were identified using Prokka (22) and clustered into orthologous gene families using Roary (16). Coinfinder took 7.2 minutes (using 20 cores; see Table 2 for more runtime details) to examine the relationships between 2,813 gene families across 534 genomes (3,957,891 pairwise tests in total). Coinfinder identified 104,944 associating gene pairs which clustered into 32 connected components or sets of genes that associate with each other. Similarly, Coinfinder took 7.5 minutes using 20 cores to identify 98,461 dissociate gene relationships within this dataset. The network and heatmap outputs of Coinfinder from this example set are shown in Figure 1.

**Table 2:** Real computational time for Coinfinder executed on a 534 genome dataset consisting of 2,813 accessory genes using different numbers of CPUs (GenuineIntel; Intel Xeon Gold 6142 CPU @ 2.60GHz)

| **Number of CPUs** | **Real computer clock time** |
| --- | --- |
| 2 | 31m16.265s |
| 4 | 17m56.973s |
| 8 | 11m15.469s |
| 16 | 7m44.942s |
| 32 | 6m16.218s |

Of the gene associations and dissociations that Coinfinder identified, many recapitulate what we know biologically. As an example, we focus on a V-ATPase present in *S. pneumoniae*. While the V-ATPase in *S. pneumoniae* has been understudied, it has been well-documented in *S. pyogenes* and sister taxon *Enterococcus hirae* (23) (24). In *E. hirae* the V-ATPase consists of 10-11 proteins organized into the ntp operon: ntpFIKECGABD(H)J (24). In *S. pneumoniae*, the V-ATPase complex is predicted to contain 9 proteins (KEGG pathway *spx_M00159*; (24)). In the annotation of *S. pneumoniae* that we performed here, only 6 genes of the ntp operon were annotated successfully: *ntpA, ntpB, ntpC, ntpD, ntpG*, and *ntpK*. Coinfinder identified consistent co-occurrence relationships between these 6 genes, forming a clique (i.e. a complete subgraph of gene associations; Figure 2a). However, these 6 genes also co-occurred with other genes in the dataset; we extended our analyses to determine whether any other genes consistently co-occurred with all 6 genes of this operon. In doing so, we identified 3 genes – atpE, and two unnamed genes – with homology to *ntpE, ntpI*, and *ntpG/H*, respectively, that consistently co-occur with the rest of the ntp operon (Figure 2a). An additional 51 genes formed cliques with the genes of the ntp operon. Of the 51 genes, 3 encode neuraminadase genes from *nan* gene clusters (Figure 2b-c). Another 3 genes co-occurring with the V-ATPase complex belong to the *dpnMAB* operon which encode the DpnII system implicated in DNA transformation (among other functions) (25) and an additional 3 are homologous to transposase IS66-related domains, perhaps suggesting how this operon has been horizontally transferred in this species (Figure 2b-c). Additionally, 4 of these proteins contained a putative cell wall binding repeat (“*CW_binding_1*”) which has been implicated in choline binding (26). Choline-binding proteins (CBPs) contain a choline-binding module/domain which allows them to bind to the cell wall of *S. pneumoniae*, functioning as essential elements of cell division, as well as strong determinants of virulence (26) (27). It is unknown why 4 CBPs co-occur with the V-ATPase complex; in eukaryotes, it has been shown that acetylcholine can be transmitted via the V-ATPase complex of vacuoles (28) but the result has not been generalized to prokaryotic cell membranes. A further 11 genes are of uncharacterised function. This example shows the power of Coinfinder in (a) identifying gene associations between proteins in a known protein complex; (b) being able to overcome poor gene annotations by looking for patterns in gene co-occurrence and gene association networks; and (c) being able to extrapolate those results to other genes with known protein interactions.

**Figure 2:**
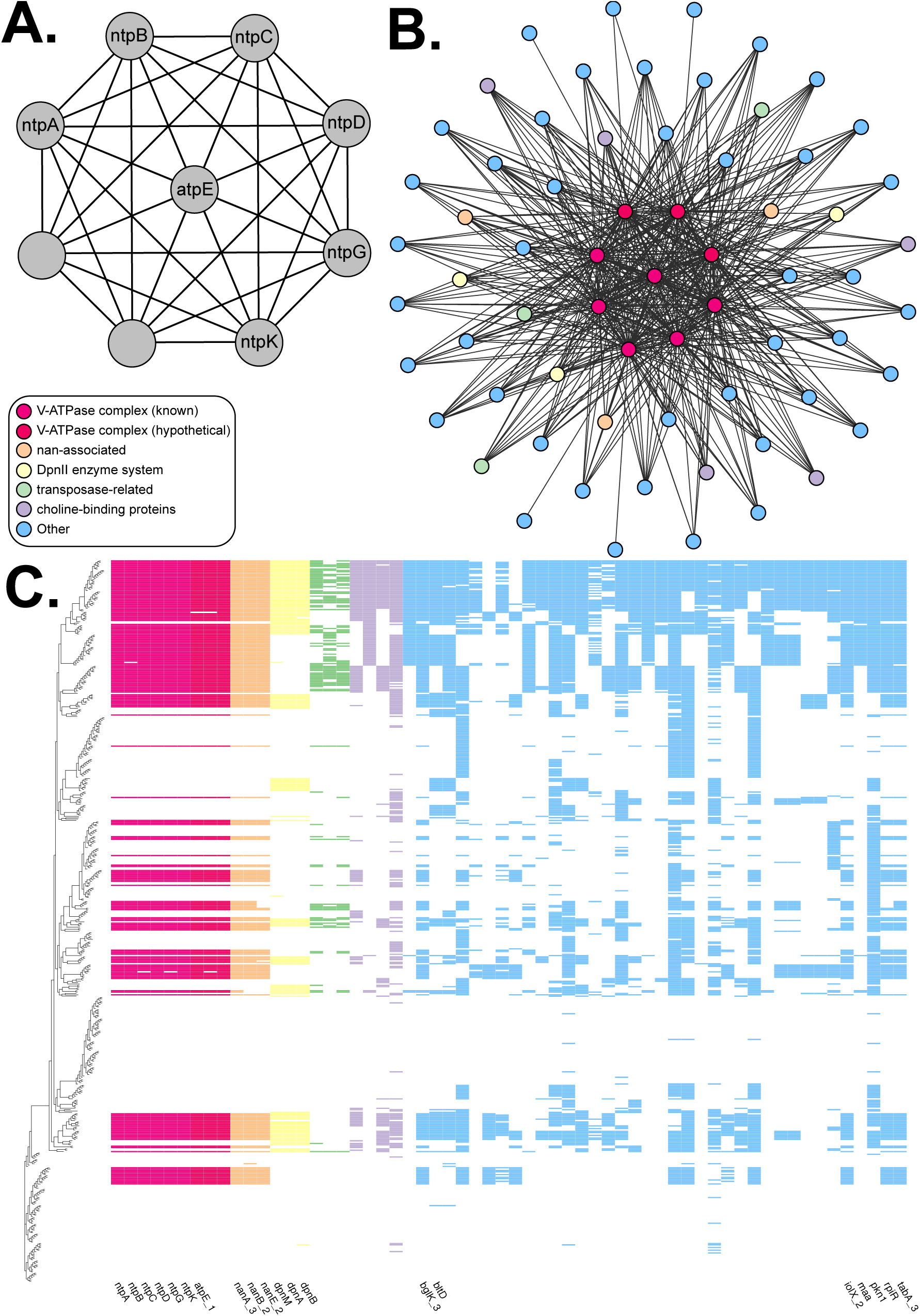
Example of the association relationships Coinfinder can identify. **A.** A clique of genes in the ntp operon which was identified within the association network (Figure 1a). 6 of these genes were correctly labelled with their gene names via the Prokka/Roary pipeline; 1 gene was given an alternative gene name often used as a synonym in the literature; a further 2 genes were listed as “hypothetical proteins”. Collectively, the 9 genes that compose the V-ATPase/ntp operon form cliques with an additional 51 genes. These cliques are shown as a network (**B**) and as a presence-absence heatmap (**C**). In the heatmap, unlabelled gene columns represent unnamed hypotheticals.

Coinfinder uses parallel processing to compute pairwise tests of coincident relationships. The most time-consuming step is the determination of the lineage-dependence of each gene; consequently, we have programmed this part to run in parallel for only those genes that are found in statistically significant coincident relationships. For the *S. pneumoniae* example, using the input set of 2,813 accessory gene families, the lineage-dependence calculation was only necessary on the 1,961 genes deemed to be in coincident relationships. Using these data, the computation time varied from 6 to 31 minutes when using 32 to 2 CPUs, respectively (Table 2).

## 8. Conclusions

Coinfinder is an accurate and efficient tool for the identification of coincident gene relationships within pangenomes. Coinfinder is open-source software available from https://github.com/fwhelan/coinfinder.

## 9. Author statements

### 9.1 Authors and contributors

FJW, MR, and JOM conceptualized this work. FJW and MR built the software. FJW validated and visualized the output data. FJW wrote the original draft; FJW, MR, and JOM reviewed and edited the manuscript. JOM acquired the funding and conducted project administration.

### 9.2 Conflicts of interest

The author(s) declare that there are no conflicts of interest.

### 9.3 Funding information

JOM was awarded funding from the BBSRC No. BB/N018044/1 to support the work of MJR and FJW. FJW has received funding from the European Union’s Horizon 2020 research and innovation programme under the Marie Skłodowska-Curie grant agreement No. 793818.

### 9.4 Ethical approval

NA.

### 9.5 Consent for publication

NA.

## 9.6 Acknowledgements

The authors would like to thank the members of the McInerney research group for valuable input, as well as the Global Pneumococcal Sequencing Project for their dedication to open-source sequencing data.

## 11. Data bibliography

1. Coinfinder is freely available at https://github.com/fwhelan/coinfinder.
2. A list of the Identifiers of the genomes used within as well as all input/output files are available https://github.com/fwhelan/coinfinder-manuscript.

## Notes

https://github.com/fwhelan/coinfinder

https://github.com/fwhelan/coinfinder-manuscript

